# GPnotebook: A pan-cancer glycoproteomic database and toolkit for analysis of protein glycosylation changes associated with cancer phenotypes

**DOI:** 10.1101/2024.04.18.589619

**Authors:** Hui Zhang, Yingwei Hu

## Abstract

Protein glycosylation plays a pivotal role in various biological processes, and the analysis of intact glycopeptides (IGPs) has emerged as a powerful approach for characterizing alterations in protein glycosylation associated with diseases. Despite the critical insights gained from IGP analysis, there is an evident scarcity of intact glycopeptide database and specialized tools for a comprehensive glycoproteomic examination. In response to this deficiency, we have developed a Python package, “GPnotebook,” which consolidates the intact glycopeptides identified from different cancer types by the Clinical Proteomic Tumor Analysis Consortium (CPTAC) and includes analytical tools for an in-depth characterization of glycopeptides. GPnotebook facilitates an array of functions including statistical profiling, differential expression analysis, glycosylation subtype categorization, investigation of glycosylation-phosphorylation interplay, survival analysis, and glycosylation enzyme assessment. We have deployed GPnotebook in a study of Pancreatic Ductal Adenocarcinoma (PDAC), thereby validating its application and demonstrating its capabilities. Our findings suggest that IGPs hold significant promise as cancer-specific changes and subtype differentiation. Consequently, GPnotebook stands out as a valuable resource for cancer researchers delving into the nuances of protein glycosylation and its correlation with cancer phenotypes.

**HILIGHTS:**

1. Simplified and unified access to pan-cancer glycoproteomic database including 90,795 intact glycopeptides.
2. Developed a glycoproteomic analysis toolkit for systematic glycoproteomic data analysis
3. Applied the toolkit for identification of glycosylation changes associated with cancer phenotypes in pancreatic cancer.

## INTRODUCTION

Protein glycosylation, the attachment of carbohydrate groups to proteins, is a vital process that plays roles in various biological functions (Pinho and Reis, 2015; Reily et al., 2019), including protein trafficking (Vagin et al., 2009), cell adhesion (Ohtsubo and Marth, 2006), immune response (Rudd et al., 2001), and receptor binding and activation (Ferreira et al., 2018). However, the analysis of glycoproteins is complex due to the non-templated nature of glycosylation, the high degree of structural diversity and isomerism among glycans, and the presence of variable occupancy of glycosites (macroheterogeneity) and multiple distinct glycan structures at each glycosite (microheterogeneity). Despite these challenges, the study of protein glycosylation is important due to its potential role in diseases and as therapeutic or diagnostic targets. Mass spectrometry, lectin binding assays, and ELISAs are among the technologies used to identify and characterize glycans and to quantify glycosylation levels at different sites on proteins (Bagdonaite et al., 2022; Goumenou et al., 2021; Pan et al., 2011; Wu et al., 2014).

Despite significant progress in the identification of intact glycopeptides using tandem mass spectrometry (Fang, Zheng et al., 2022; Polasky et al., 2020; Shen et al., 2021; Toghi Eshghi et al., 2015; Zeng et al., 2021) and glycan databases (Alocci et al., 2019; Fujita et al., 2021; Minoru Kanehisa, 2017), the high-throughput analysis of glycoproteomic data still faces considerable obstacles. A prominent challenge is the lack of specialized resources tailored to the glycoproteomic evaluation of intact glycopeptides (IGPs). This deficiency hinders the seamless integration of glycoproteomic information with other ‘omics’ datasets. Furthermore, there is a void in the availability of tools specifically calibrated for glycoproteomic analysis that can robustly link identified glycosylation alterations to cancer phenotypes, with an emphasis on reproducibility for subsequent validation and research. Addressing these challenges is paramount to unlock the full potential of glycoproteomics in cancer and the broader understanding of disease mechanisms.

We presented here a glycoproteomic analysis toolkit, GPnotebook, as a promising approach to high-throughput glycoproteomic data analysis for IGPs. Using the mass spectrometry data sets from 10 tumor types characterized by CPTAC projects, we created a python package, named as ‘cptac_glyco’, to provide the unified access to identified glycopeptides of CPTAC cancer glycoproteomics data, containing 90,661 N-linked glycopeptides from 2,194 proteins. The 10 cancer types include breast carcinoma (BRC) (Krug et al., 2020), clear cell renal cell carcinoma (ccRCC) (Clark et al., 2019), colorectal carcinoma (CRC) (Vasaikar et al., 2019), glioblastoma (GBM) (Wang, L. et al., 2021), head and neck squamous cell carcinoma (HNSCC) (Huang et al., 2021), lung squamous cell carcinoma (LSCC) (Satpathy et al., 2021), lung adenocarcinoma (LUAD) (Gillette et al., 2020), ovarian serous cystadenocarcinoma (OVC) (Hu et al., 2020), pancreatic ductal adenocarcinoma (PDAC) (Cao et al., 2021), and uterine corpus endometrial carcinoma (UCEC) (Dou et al., 2020). Based on the standardized expression matrices, we developed GPnotebook for the comprehensive glycoproteomic data analysis based on IGPs. The performance of the data analysis tools demonstrated through its applications in statistical profiling, differential expression analysis, glycosylation subtype categorization, investigation of glycosylation-phosphorylation interplay, survival analysis, and glycosylation enzyme assessment for a PDAC cohort.

## RESULTS

### Landscape of the glycoproteomic database

Since phosphoproteomic data set include a large portion of glycoproteomic spectra, especially IGP with sialic acids are co-enriched with phosphopeptides (Hu et al., 2018). The data sources of both glycoproteomic and phosphoproteomic data sets of tumor and normal adjacent samples of the 10 CPTAC cohorts were used for the assignment of IGPs (Figure 1A). The corresponding clinical information, including stage, grade, and overall survival time was also collected from the literatures (Figure 1B). Our goal was to search for intact glycopeptides, and in this initial study, we reported 90,661 glycopeptides in the database under a glycopeptide-spectrum matching (GPSM) false discovery rate (FDR) < 0.01 threshold (Figure 1C).

**Figure 1.**
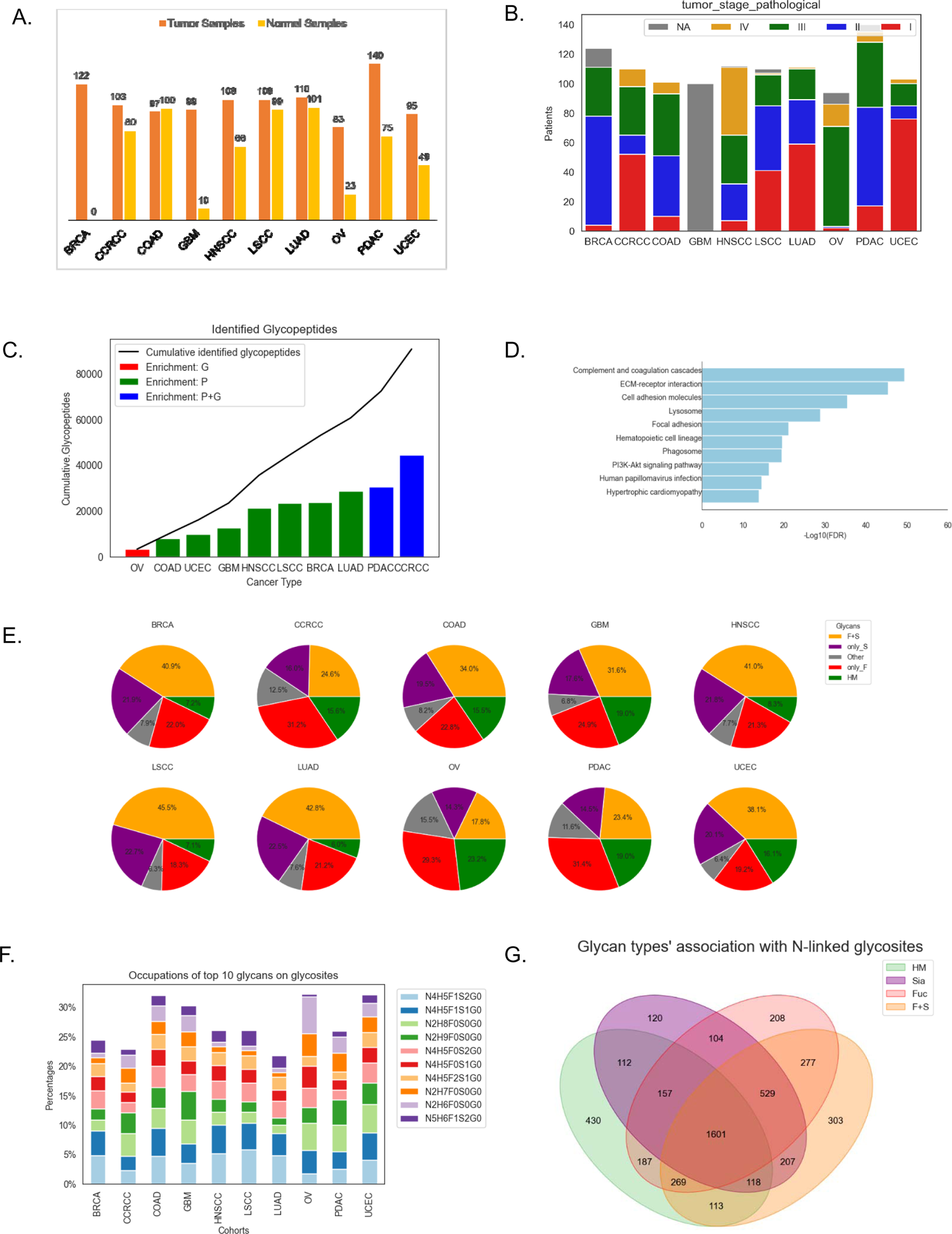
Landscape of the glycoproteomic database (A) Statistics of patients, tumor, and non-tumor samples (B) Clinical information (stage) of patients in all the 10 cohorts (C) Statistics of intact glycopeptide identifications from the 10 cohorts (D) KEGG pathways enriched from the identified glycoproteins (E) Glycans identified in the 10 cohorts (F) The glycans that are most frequently identified in each cancer type (G) The association of glycan types with glycosites across the 10 cancer types

To gain insights into which of cellular components were included in the identified glycoproteins, we conducted enrichment analysis of all the identified glycopeptides using GSEApy (Fang, Zhuoqing et al., 2023) and KEGG pathway database (Kanehisa, M. and Goto, 2000; Kanehisa, Minoru et al., 2016). Our results showed that the top three enriched pathways were complement and coagulation cascades, ECM-receptor interaction, and cell adhesion molecules, all under an FDR < 0.05 threshold (Figure 1D).

We also characterized the glycans associated with the identified glycopeptides in the database for each cohort (Figure 1E). Our results indicated that the glycan proportions varied across different cancer types, likely due to both the use of different glycoproteomic analysis techniques and sample types. However, the most common glycan types were consistent across all 10 CPTAC cohorts. Specifically, oligomannose (N2H6-N2H9) and N4H5-cored complex glycans were the most common N-linked glycans attached to the identified glycopeptides, occurring on approximately 25% of the total glycopeptides on average (Figure 1F). Furthermore, we investigated the associations between glycosites and glycans and found that approximately 30% of glycosites were modified by different N-linked glycans (Figure 1G). This finding is useful for studying the microheterogeneity of glycans on glycosites.

### Differentially expression of N-linked glycopeptides between Tumor and NAT samples in PDAC

Differentially expressed glycopeptides (DEGPs) from each cancer types can be identified from the database. We integrated OmicsOne to establish a pipeline to identify DEGPs between tumor and NAT (normal adjacent tissue) samples in various datasets, characterize the properties of DEGPs, and investigate the association between clinical information and DEGPs (Zhang et al., 2021). Pancreatic cancer is still a leading cause of death globally, and early detection is crucial for improving survival rates. Our results, depicted in Figure 2A, indicate that the differential expression of glycopeptides can effectively distinguish tumor and NAT samples in most cases.

**Figure 2.**
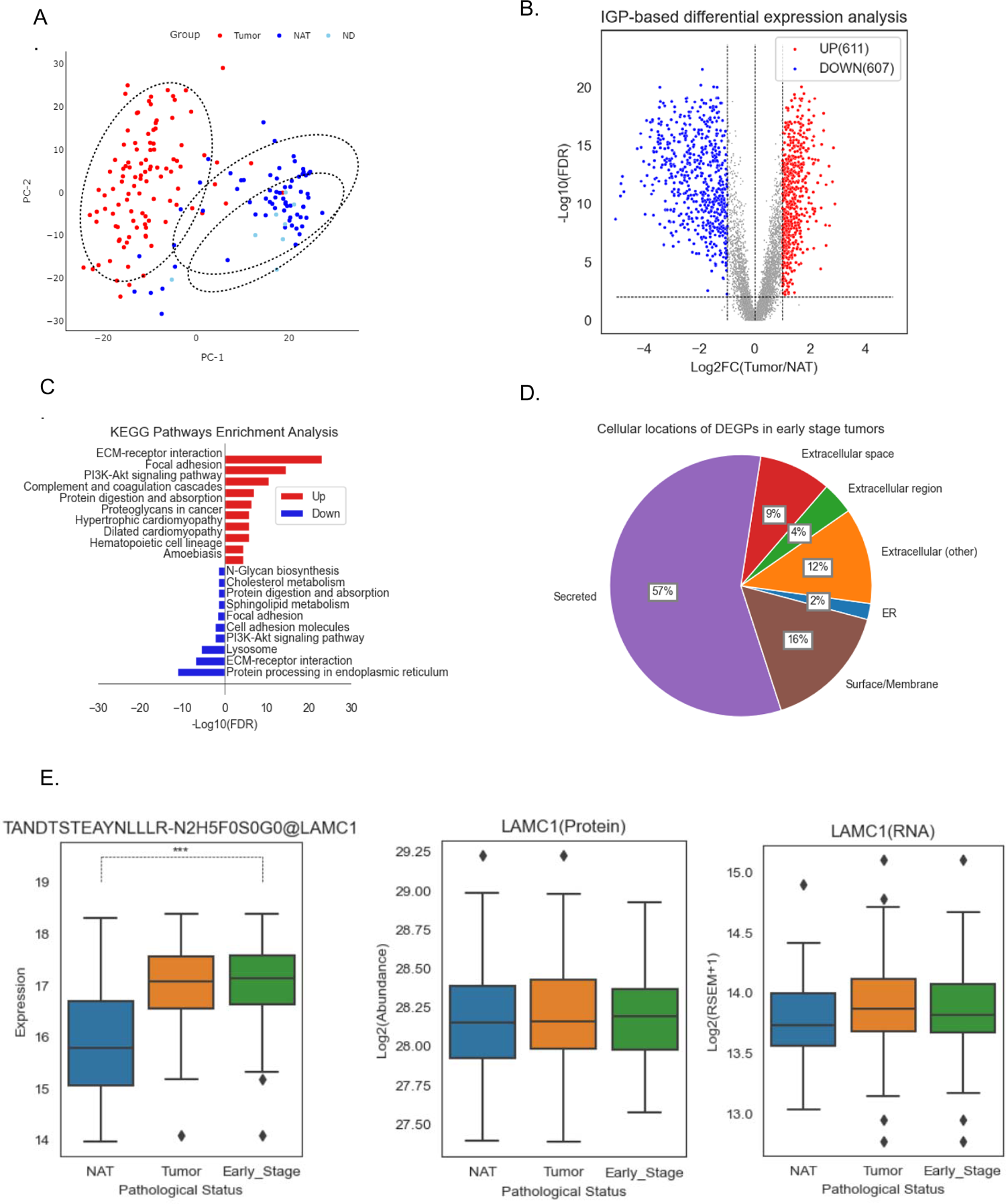
Differentially expression of N-linked glycopeptides between Tumor and NAT samples in PDAC (A) Principal component analysis of tumor, normal adjacent (NAT), and normal ductal tissues (B) Volcano plot of differentially expressed intact glycopeptides (IGPs) in tumor comparing to NATs (C) KEGG Pathways enriched in up-regulated and down-regulated IGPs (D) The cellular locations of differential expressed glycopeptides in early stage tumors (E) Multi-omics expression profiles of TANDTSTEAYNLLLR-N2H7F0S0G0@LAMC1 in NATs, tumors, and early stage tumors, including IGP (left), proteomic (middle), and transcriptomic (right).

We identified 1,218 DEGPs corresponding to 260 glycoproteins in the comparison between 105 PDAC samples and 67 NAT samples (as well as 8 normal ductal samples (ND)) from the PDAC dataset using the Wilcoxon rank sum test followed by multiple testing correction (FDR <0.01), of which 607 are significantly down-regulated, and 611 are significantly up-regulated (fold change > 2) (Figure 2B). The enriched pathways in up-regulated and down-regulated glycopeptides are illustrated in Figure 2C, with ECM-receptor interaction, focal adhesion, and PI3K-Akt signaling pathway being the top three pathways enriched in the gene names of significantly up-regulated glycopeptides. In contrast, protein processing in endoplasmic reticulum, ECM-receptor interaction, and lysosome are the top three pathways enriched in the gene names of significantly down-regulated glycopeptides.

We further screened for glycopeptides associated with early stage of PDAC by comparing 83 early-stage (stage I and II) tumor samples with NAT samples and found 366 glycopeptides (corresponding to 101 glycoproteins) with significant up-regulation. Out of the 101 glycoproteins, 57% (83 glycoproteins) were related to secreted or extracellular proteins (Figure 2D). Comparing the results with corresponding proteomic data for global protein expression, we found that 50 out of the 83 proteins were not significantly up-regulated (fold change > 2) at the total protein level. Our findings suggest that glycopeptide expression can provide new insights for early detection compared to protein expression data. As an example, the expression profiles (including IGP, protein, and RNA) of a sample glycopeptide TANDTSTEAYNLLLR-N2H7F0S0G0@LAMC1 in NATs, tumors, and early-stage tumors were presented in Figure 2E.

### Glycoproteomics based subtyping of PDAC

A study was conducted to investigate tumor heterogeneity in PDAC using differential expression data of glycopeptides (IGPs) to cluster 105 tumor samples. Non-negative matrix factorization (NMF) clustering (Gaujoux and Seoighe, 2010) was employed, and three tumor subtypes were identified based on glycoproteomics data. The study subsequently integrated multi-omics-based NMF subtypes (C1 and C2) of PDAC previously annotated by CPTAC, *KRAS* mutation annotation, and xCell (Aran et al., 2017) immune clustering results (Figure 3A). Enrichment scores were evaluated between different omics data types to investigate the relationship between glyco-based subtypes and other molecular-based classifications of PDAC (Cao et al., 2021), including tumor cellularity, different *KRAS* mutations, immune cell cluster, and multi-omics NMF clustering. A hypergeometric test p value of 0.05 was selected as the threshold for overlap between different classification results. The study found that among the 105 PDAC samples, samples with *KRAS*.G12V mutation were enriched in IGP cluster 3. The results also showed that the IGP-based IGP 1 and IGP2 exhibit significant overlaps with the C1 subtype, while subtype3 was consistent with the C2 cluster using multi-omics data (Figure 3B). Furthermore, as illustrated in Figure 3C, the IGP1 displays an overlap with immune cluster A (acinar-high), whereas the IGP3 exhibits an overlap with immune cluster B (immune-cold). The IGP2 demonstrates a predominant overlap with immune cluster C (immune-cold), followed by the immune cluster D (immune-hot). Glycoproteins are mainly secreted or membrane proteins involved in the immune response, and the fucosylated and sialylated glycopeptides in cluster IGP2 were relatively up-regulated compared with the corresponding glycopeptides in IGP1 and IGP3 (Figure 3D), suggesting that the samples of different IGP-based subtypes may have different immune responses. Survival analysis was also performed for the three clusters defined by the differential expression of IGPs, and a Kaplan-Meier plot showed that IGP3 subtype had the worst 3-year survival prognosis, while IGP 2 subtype had the best one (Figure 3E). The study suggests that the signature glycopeptides contributed to construct the IGP2 could be useful indicators in prognosis.

**Figure 3.**
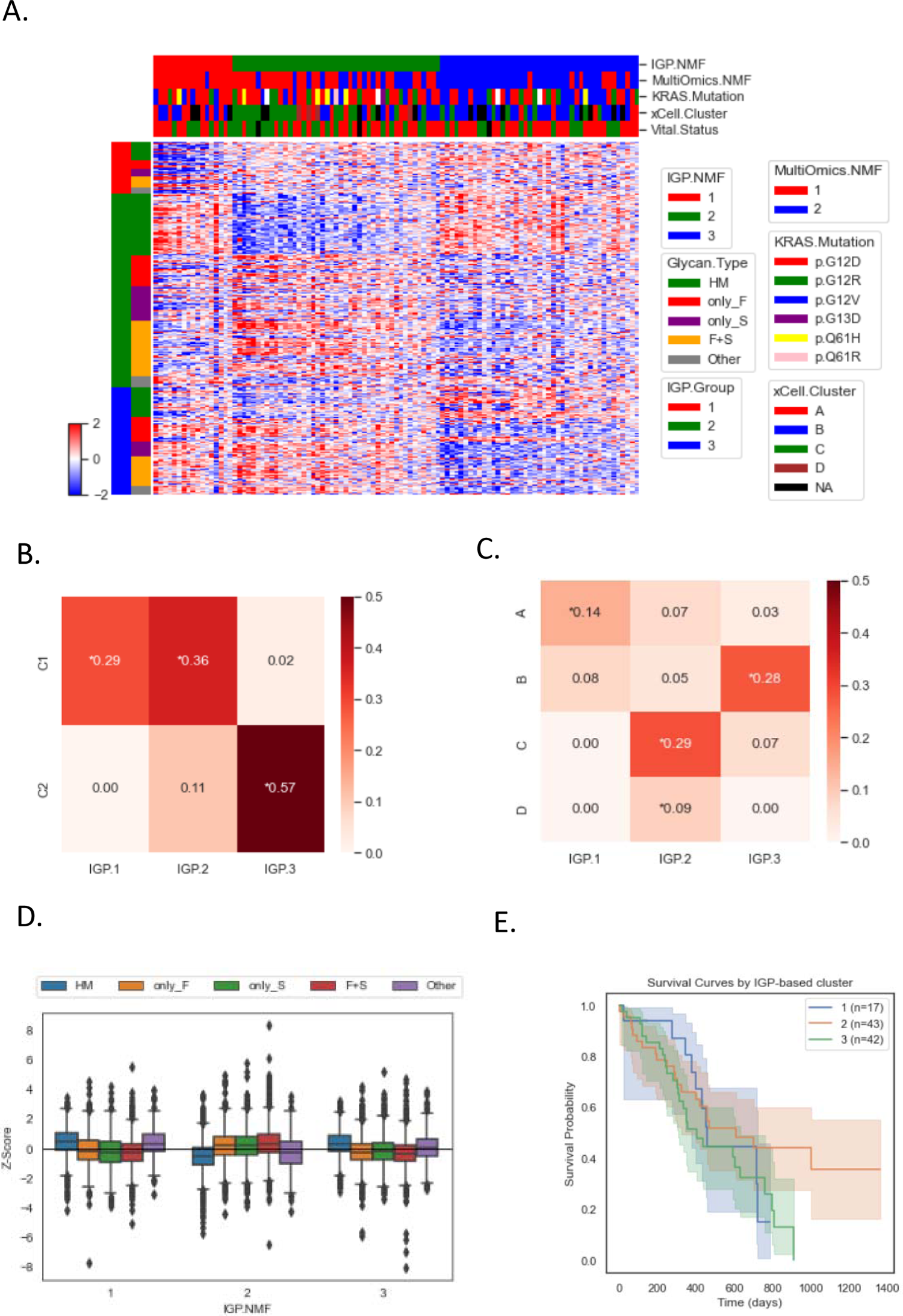
Glycoproteomics based clustering of PDAC (A) IGP-based NMF clustering (B) Comparison between IGP-based and multiomics-based clustering results (C) Comparison between IGP-based clusters and xCell immune clusters (D) Differential glycan expression in clusters (E) Survival analysis of IGP-based clusters

### Survival-associated glycopeptide expression signature in PDAC

We collected data from 105 tumor patients and selected 85 cases, including 42 living patients and 43 patients whose cause of death is PDAC, with a maximum follow-up period of 1364 days (Figure 4A). A survival analysis pipeline was established in GPnotebook using lifelines (Davidson-Pilon, 2019) and scikit-survival (S. Pölsterl, 2020) to analyze the associations between overall patient survival and intact glycopeptides expression. The Kaplan-Meier curve of overall survival (OS) time was presented in Figure 4B, which demonstrated that the survival probability declined linearly at a rate of about 30% annually in the first three years. Through univariate Cox regression analysis in lifelines, as shown in Figure 4C, we displayed the top 20 IGPs that were significantly associated with the OS of PDAC cancer patients (adjusted p-value < 0.05). Among these, the three intact glycopeptides of TNC and one intact glycopeptides of LAMA4 had the highest hazard ratio values. The time-dependent risk analysis of these three intact glycopeptides of TNC revealed similar trends of high risks (threshold >0.5) during the time range of tumor development, particularly in the first year (Figure 4D). We employed a penalized COX model with elastic net penalty score for further feature selection and constructed a multivariate COX regression model using 8 selected prognostic IGPs optimized from 80% training set of the data (N=68). The coefficient values of these 8 IGPs in the model were presented in Figure 4E. Lastly, we performed the multivariate COX regression model on the remaining 20% testing data (N=17) and found that the model exhibited moderate performance with an median AUC of nearly 0.7 (Figure 4F). The result demonstrates that the IGP-based COX model can effectively predict survival chance in the PDAC cohort.

**Figure 4.**
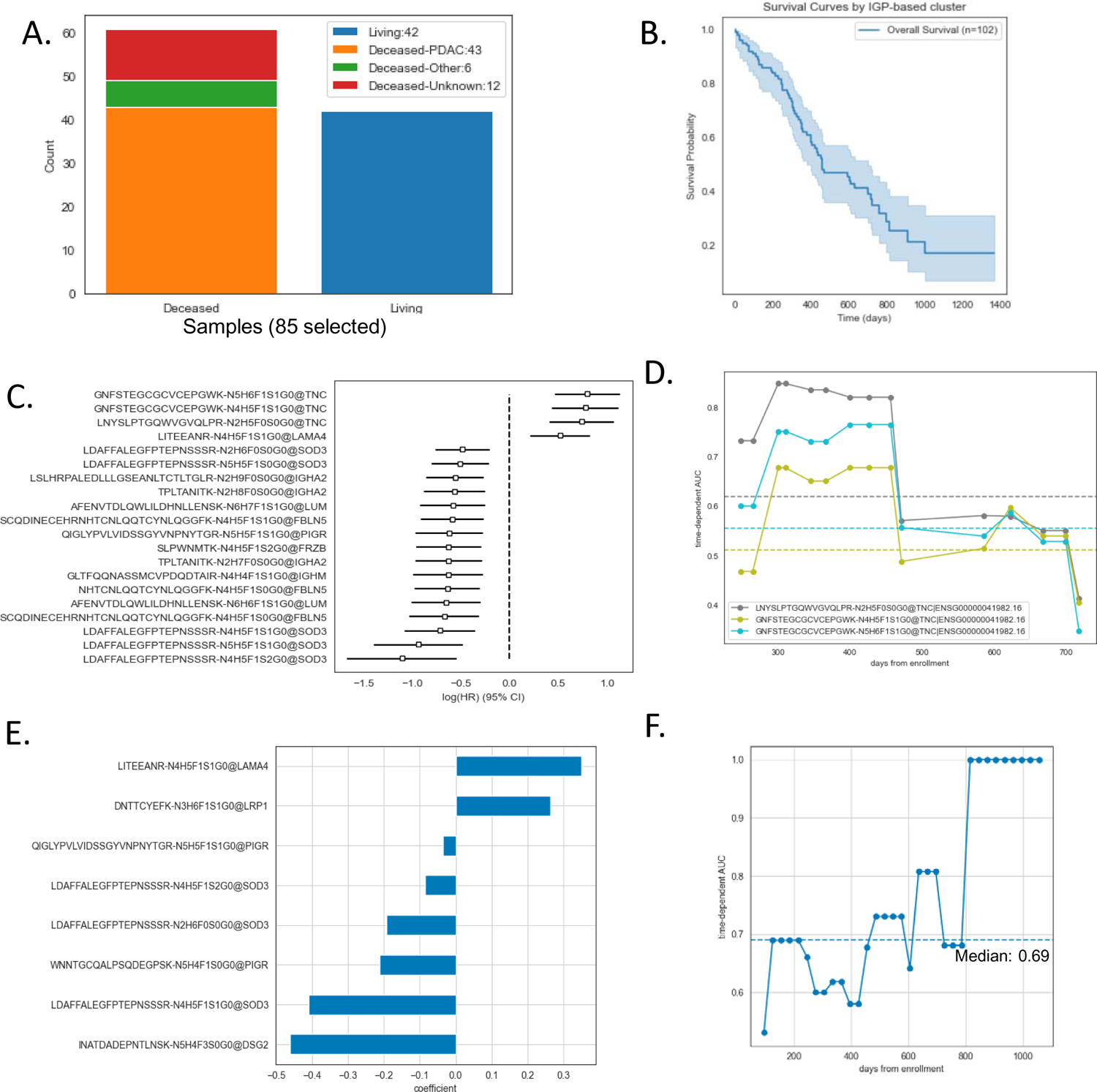
Survival-associated glycopeptide signatures in PDAC (A) Survival status of patients (B) The Kaplan-Meier (K-M) curve of over all survival time of 102 patients (C) Hazard ratio estimation of univariate models (D) Time-dependent area curves under the ROC of the univariate models based on the specific glycopeptides of TNC (E) The coefficients of multivariate variables in the COX model, which was penalized by the elastic net method. (F) Time-dependent area under the ROC of the multivariate COX model estimated on testing data

### Glycosylation-phosphorylation cross-talk in PDAC

Phosphorylation is known to play crucial roles in the regulation of various signaling pathways (Ardito et al., 2017). However, there are reports that signaling pathways could be regulated by protein glycosylation (Wang, X. et al., 2006). To investigate the regulation of glycosylation and phosphorylation, we further aimed to identify signaling pathways affected by glycosylation aberrations. Specifically, we examined two conditions: intra- and inter-cross talk between glycosylation and phosphorylation. The former refers to the proteins that have identifications of both phosphorylated and glycosylated peptides. The latter refers to correlation between all identified glycopeptides and phosphopeptides from all proteins.

Among 1,123 glycoproteins and 7,733 phosphoproteins analyzed in 105 PDAC samples, we found that 502 proteins had both glycosylation and phosphorylation sites. Notably, more than half of these glycoproteins were membrane proteins, according to their cellular component gene ontology annotations. The overrepresentation analysis using GSEApy on hallmark pathway database revealed that pathways of epithelial mesenchymal transition, coagulation, myogenesis, apical junction and angiogenesis were enriched in the 502 intra-cross talk proteins (Figure 5B). We also found that the expression values of glycopeptides and phosphopeptides of the identical proteins were positively proportional (p value < 0.01) (Figure 5C), although the phosphorylation and glycosylation modifications on the same protein are not always strongly correlated. For instance, the phosphosite S1222 of LAMB1 was significantly down-regulated, while the two glycopeptides (GTNYTVR-N2H7F0S0G0 and LSDTTSQSNSTAK-N3H3F1S0G0) of LAMB1 were significantly up-regulated. Conversely, the phosphopeptides of HSPG2, SUN2, and SOD3 were significantly up-regulated while their glycopeptides were significantly down-regulated. The inconsistent changes in glycosylation and phosphorylation indicate the complexity and diversity of protein modification regulation.

**Figure 5.**
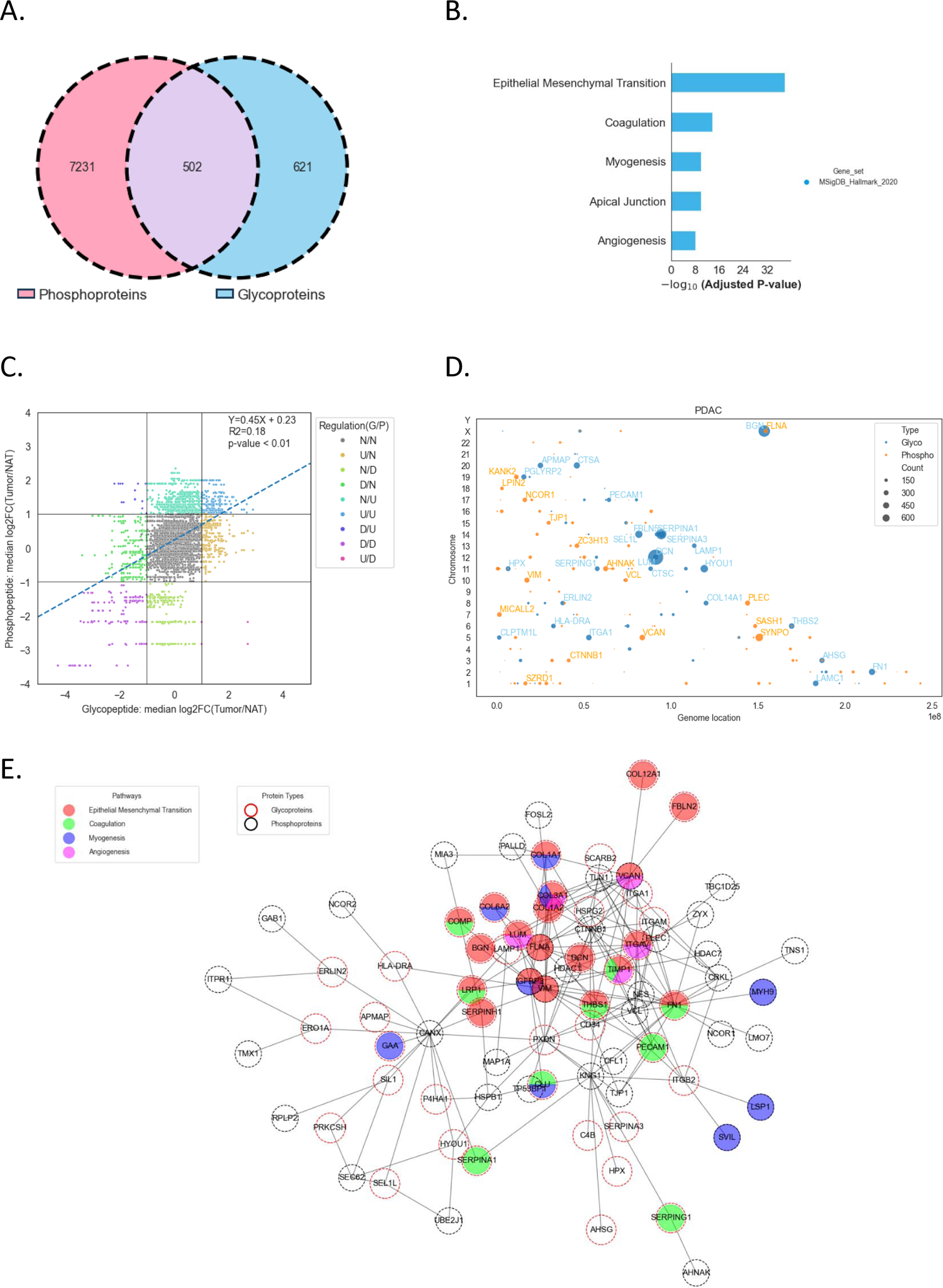
Glycosylation-phosphorylation cross talk in PDAC (A) Venn diagram of glycoproteins, phosphoproteins, and phospho-glycoproteins (B) Hallmark pathways enriched in phospho-glycoproteins (C) Differential expression of phosphopeptides and glycopeptides from phospho-glycoproteins (D) The chromosome locations of glycopeptides (blue) and phosphopeptides (orange) which significantly altered in tumor tissues compared to NATs and exhibit significant highly positive spearman correlations (r > 0.6) with each other. The numbers of high correlations were highlighted by marker sizes. (E) Protein-protein interaction network between positively correlated glycoproteins and phosphoproteins (corr.r > 0.6 and corr.size > 10).

In order to further explore the relationship between protein glycosylation and phosphorylation across different proteins, additional analyses were conducted. The inter-cross talk between protein glycosylation and protein phosphorylation expressed in all the 105 PDAC tumor samples were studied by GPnotebook. Figure 5D displays the genomic locations of glycopeptides and phosphopeptides that demonstrate at least 10 positive correlations (spearman.corr > 0.6) with each other, provided that either the glycopeptide or the phosphopeptide is significantly up- or down-regulated in the correlation. DCN, BGN, SERPINA1, HYOU1, and SERPINA3 are the top 5 gene names of glycopeptides that have the most positive correlations with phosphopeptides. SYNPO, VCAN, VIM, PLEC, and AHNAK are the top 5 gene names of phosphopeptides having the most positive correlations with glycopeptides. The over-representation analysis of all phosphopeptides correlated with glycopeptides of BGN demonstrated that hallmark pathways of Mitotic Spindle, Epithelial Mesenchymal Transition, and apoptosis are enriched in the gene names of positively correlated phosphopeptides (FDR < 0.05).

The protein-protein interaction network analysis of the gene names of strongly correlated glycopeptides and phosphopeptides (> 10 positive correlations and spearman corr > 0.6) using the API functions of STRINGdb (Szklarczyk et al., 2021) showed the evidence connections between glycoproteins and phosphoproteins. The integrated enrichment analysis using GSEApy further revealed that hallmark pathways of Epithelial Mesenchymal Transition, Coagulation, Myogenesis, and Angiogenesis are the main pathways involved in the glycoprotein-phosphoprotein interaction network, as shown in Figure 5E, shedding new light on the regulation of protein modification in pancreatic cancer study.

### Impact of glycogene expression on protein glycosylation in pan-cancer studies

The regulation of protein glycosylation involves a complex network of enzymes that facilitate the attachment and removal of specific carbohydrate molecules from proteins. The process of attaching carbohydrate molecules to proteins is mediated by glycosyltransferases, while the process of hydrolyzing glycosidic bonds between carbohydrates and other molecules is catalyzed by glycosidases. In this study, we investigated the expression of 111 enzymes involved in protein glycosylation biosynthesis (http://glycoenzymes.ccrc.uga.edu/Glycomics3/) in the 10 CPTAC cohorts.

Using RNA-seq data, we were able to identify all 111 enzymes at the transcriptome level in most of the CPTAC cohorts, with the exception of colorectal cancer (CRC), glioblastoma (GBM), and ovarian cancer (OVC). More than half of the enzymes were also identified in the proteomic data (Figure 6A). Our analysis revealed that the protein expression of these enzymes is related to their RNA expression, but also regulated by other factors, as indicated by median correlation values ranging from 0.25 to 0.5 (Figure 6B).

**Figure 6.**
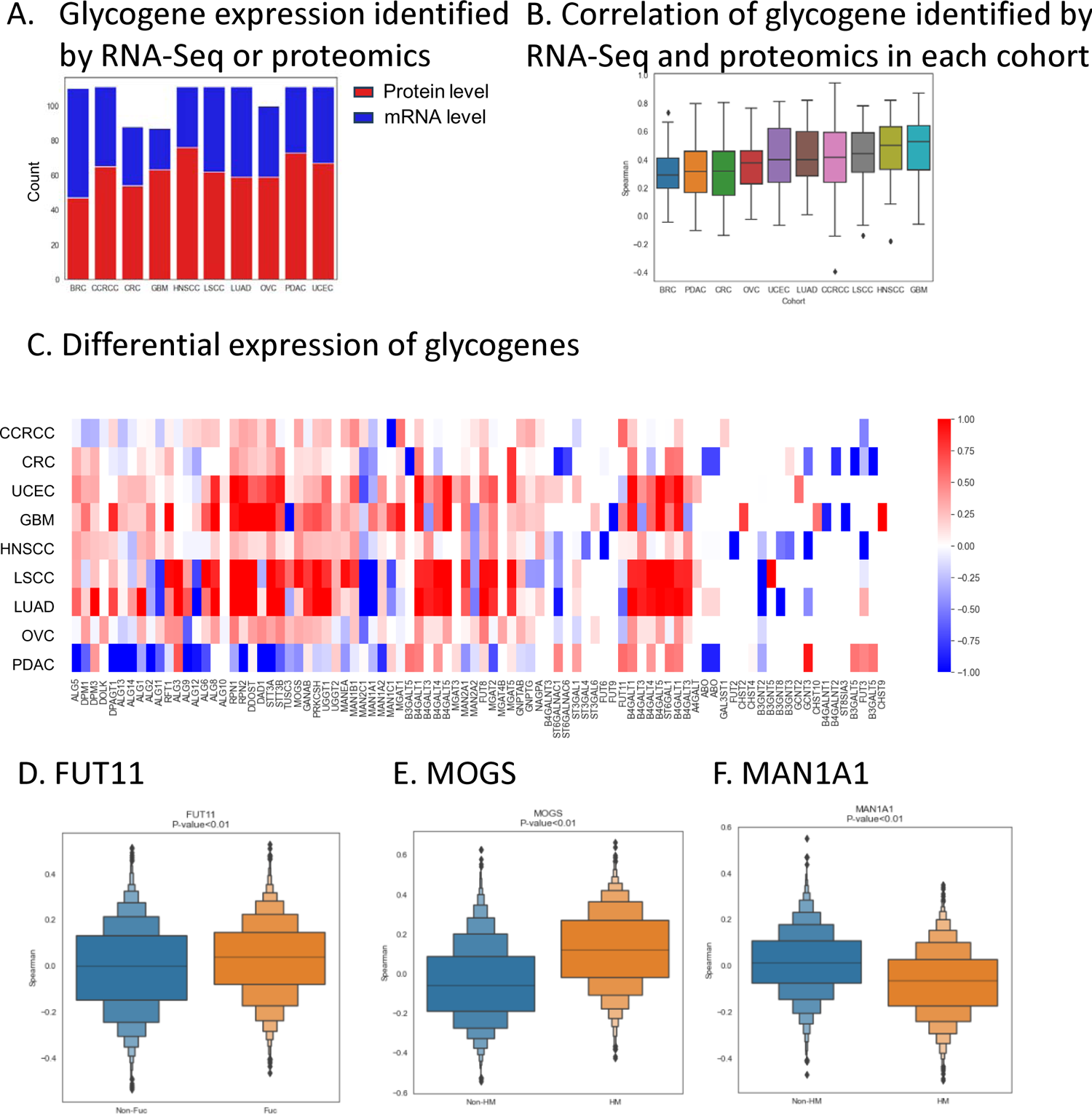
Impact of glycogene on protein glycosylation in pan-cancer studies (A) Identification of enzymes involved in biosynthetic glycosylation across 10 cancer types. The bar chart depicts the enzymes discovered through RNA-Seq (blue) and proteomic (red) experiments (B) Correlation of enzymes identified by RNA-Seq and proteomics in each cohort (C) Heatmap illustrating the differential expression of enzymes in various cancer types. The color-coded matrix presents the fold change in protein expression levels of enzymes between tumor samples and their corresponding normal adjacent tissues (NATs). Upregulated enzymes are represented in red, while downregulated enzymes are shown in blue, indicating the heterogeneity in proteomic profiles across different cancer types. (D) Correlation between FUT11 and IGPs with/without Fuc glycans (Fuc and non-Fuc). (E) Correlation between MOGS and IGPs with/without Oligomannose glycans (HM and non-HM). (F) Correlation between MAN1A1 and IGPs with/without Oligomannose glycans (HM and non-HM).

Furthermore, we observed that most of the glycosylation enzymes in the precursor pathway are up-regulated in tumor samples compared to normal adjacent tissue (NAT) samples, except for PDAC. In addition, we found that MAN1A1 and MAN2C1 are down-regulated in most cancer types, while B4GALT1, and B4GALT3-5 are up-regulated in most cancer types (Figure 6C). These findings suggest that dysregulation of glycosylation enzymes may contribute to tumorigenesis.

Lastly, our analysis revealed significant differences in the correlation between different glycan types and certain glycosylation enzymes. For example, in the PDAC cohort, FUT11 had a higher positive correlation with fucosylated glycopeptides (Figure 6D), while MOGS had a higher positive correlation with oligomannosylated glycopeptides (Figure 6E). Conversely, MAN1A1 had a negative correlation with oligomannosylated glycopeptides (Figure 6F). The findings suggest that modifications to the expression of glycosylation enzymes have an effect on the production of glycoproteins’ end products in the PDAC tumor samples.

### GPnotebook for glycoproteomic analysis in cancer studies

We developed GPnotebook, a toolkit that seamlessly integrates multiple data analysis workflows for comprehensive glycoproteomic investigation in cancer studies (Figure 7A). In this workflow, the intact glycopeptides were characterized by gene name, sequence, glycosite, and glycan composition. The meticulous dissection of these components is helpful to elucidating the glycopeptide’s biological roles and implications in disease pathology from multiple perspectives.

**Figure 7.**
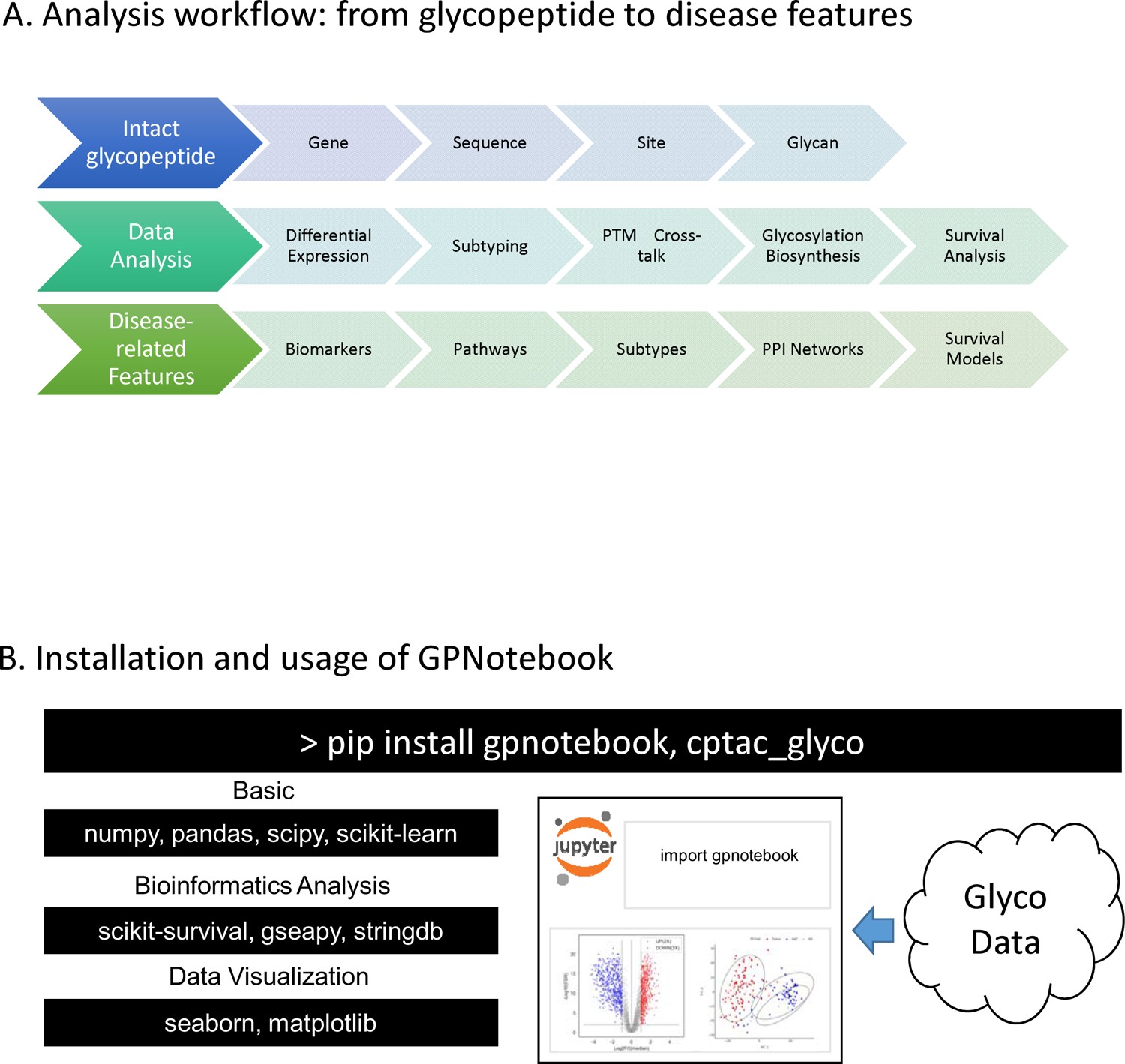
Active database for glycoproteomic in cancer studies. (A) Workflow of GPnotebook: from glycopeptides to disease features (B) Installation and infrastructure of GPnotebook

Advancing into the realm of data analysis, we apply rigorous data analysis methodologies, such as differential expression analysis, subtyping analysis, glycosylation-phosphorylation cross talk, glycosylation biosynthesis analysis, and survival analysis. The zenith of this analytical journey is the distillation of disease-relevant markers, comprising the identification of biomarkers and pathway enrichment through differential expression studies, the construction of molecular subtype classifications via IGP-based cluster analysis, the mapping of protein interaction networks informed by PTM interplay, the regulatory mechanisms of glycosylation enzymes as deciphered from biosynthetic pathway evaluations, and the development of IGP-informed prognostic models. This comprehensive methodology embodies a new standard for molecular disease research, promoting the evolution of diagnostic precision and the tailoring of therapeutic strategies. GPnotebook can be installed by the standard pip install approach (Figure 7B). It integrated several existed tools to construct the analysis pipelines for different purposes.

The basic level of the software includes numpy (numeric calculation), pandas (data table operation), scipy (statistics calculation), and scikit-learn (machine learning). The bioinformatics analysis includes in-house codes and three main packages of scikit-survival (survival analysis), gseapy (enrichment analysis), and stringdb (PPI analysis). The results were exported as tables and figures generated by seaborn and matplotlib. The visualized data were displayed in Jupyter notebook with the codes using the glycopeptide expression matrices fetched from cloud storage.

## DISCUSSION

GPnotebook is a novel glycoproteomic database and toolkit designed for intact glycopeptide analysis in cancer studies. The primary focus of the GPnotebook is to aid in the discovery and interpretation of disease-associated glycosites, intact glycopeptides, and glycoproteins, thereby complementing existing cancer data portals and glycoproteome data portals. Using the datasets from CPTAC studies and the database tools developed for the database, we conducted five case studies (Figure 2-6) of intact N-linked glycopeptide analysis to demonstrate the practical utility of GPnotebook and the potential of glycopeptides as essential indicators in the diagnosis, prognosis, or network of protein glycosylation in tumor samples.

To investigate the potential role of cancer-associated glycopeptides in the diagnosis and prognosis of PDAC, we performed differential expression analysis, tumor subtyping analysis, and survival analysis. In the differential expression analysis case study, we revealed that the differential expression of intact glycopeptides could shed new light on the protein alteration in tumor tissues. For an example, LAMC1 is a crucial subunit of the laminin gene family, non-collagenous glycoproteins found in basement membranes, and studies have linked abnormal expression of these genes to biological and clinical features of cancers, such as gastric cancer (Bizama et al., 2014), hepatocellular carcinoma (Jhunjhunwala et al., 2014), renal cell carcinoma (Wragg et al., 2016), colorectal cancer (Bartolini et al., 2016), and lung cancer (Sathyanarayana et al., 2003). Although no significant alterations were observed on the transcriptomic and proteomic levels of LAMC1 in our cohort, the significant up-regulation was only observed in the glycoproteomic data. This finding emphasizes that studying protein glycosylation is a critical complement in omics studies of cancers. In the tumor subtyping case study, we demonstrated that IGP-based clustering was related to the immune types of the tumor samples. The signature glycopeptides that contributed to the clustering may associate with their immune responses and influence the survival probability. In the survival analysis case study, we set up a pipeline to realize a multivariate regression analysis on high-dimensional data containing highly correlated features. The trained regression model showed moderate performance for the time-dependent survival probability prediction and illustrated the significant survival association with the expression of N-linked glycopeptides of LAMA4, LRP1, PIGR, DSG2, and SOD3. SOD3, short for Superoxide Dismutase 3, is an antioxidant enzyme and plays a crucial role in the body’s defense system against oxidative stress. It is interesting to see that the different glycosylation types on the same glycosite of SOD3 all benefit the better prognosis but have different effects, which may indicate that different glycosylation may influence the enzyme functions to protect cells. However, the limited samples may still influence the model performance, especially for the medium-term (200-400 days) survival prediction.

To investigate the regulation network of protein glycosylation, we performed PTM cross-talk analysis and glycogene analysis. In the PTM cross-talk case study, we provided a comprehensive analysis of protein glycosylation and phosphorylation cross-talk. We found that protein glycosylation and phosphorylation are highly associated in PDAC. About half of the identified glycoproteins have phosphorylation sites. The inconsistency of the alterations of protein glycosylation and phosphorylation indicated that protein modification can be regulated in multiple ways. We also found a large number of highly correlated glycopeptides and phosphopeptides. Referring to the interaction database, we showed a protein-protein interaction network for the most positively correlated glycoproteins and phosphoproteins. The network revealed that the glycosylation alterations in tumor samples broadcast from cells’ inside (protein processing in ER) to outside (ECM-receptor interaction). Certain phosphoproteins show the potential to play critical roles in propagating glycosylation changes. In the case study of glycosylation biosynthesis, we investigated the enzymes involved in the glycosylation biosynthesis in all the 10 CPTAC cancer cohorts from both transcriptomic and proteomic levels. The changes of glycosylation enzymes indicate that different tumor cells should have different immune responses because most of the influenced glycoproteins are secreted proteins or membrane proteins involved in immune responses.

In summary, we established a comprehensive data analysis pipeline for intact glycopeptide analysis in cancer studies by integrating the most widely-used and well-understood approaches in biomedical research. The notebooks in the GPnotebook make it easy for users to reproduce the analysis and understand the results. Although the current version of GPnotebook includes only CPTAC data, it can be easily extended to support other cohort-based glycoproteomics studies.

## START METHODS

### Data source

GPnotebook extends the CPTAC database and added glycoproteomic expression matrices for primary tumors and normal adjacent samples from the glycoproteomic 10 CPTAC cancer types from a total 1,119 patients (Figure 1A). GPnotebook provides the tools for comprehensive glycoproteomic analysis and integrates the glycoproteomic information with transcriptomic, proteomic, phosphoproteomic, and clinical data for the 10 CPTAC cancer types downloaded from the python package, CPTAC (Lindgren et al., 2021), and literatures (Cao et al., 2021; Clark et al., 2019; Dou et al., 2020; Gillette et al., 2020; Hu et al., 2020; Huang et al., 2021; Krug et al., 2020; Satpathy et al., 2021; Vasaikar et al., 2019; Wang, L. et al., 2021). The clinical data involved in this database includes tumor stage, gender, age, tumor site, vital status, and overall survival time. Substage level data were merged under respective stage (I, II, III, and IV, See Figure 1B). The molecular subtype annotation and tumor purity information were obtained from literature. All molecular data were properly normalized and stored as feature by sample matrix files.

### Database construction and characterization

The identification of intact N-linked glycopeptides in this study was carried out using GPQuest 2.1 software (Hu et al., 2018; Mertins et al., 2018). GPQuest 2.1 was used to identify intact glycopeptides from MS/MS spectra using two approaches: searching for spectra containing oxonium ions (“oxo-spectra”) and identifying intact N-linked glycopeptides.

The oxonium ions were used as signature features of glycopeptides from MS/MS spectra, which were caused by the fragmentation of glycans attached to intact glycopeptides in the mass spectrometer. In this study, the MS/MS spectra containing oxonium ions (m/z 204.0966) in the top 10 abundant peaks after removing TMT reporter ions were considered as potential glycopeptide candidates. The intact N-linked glycopeptides were identified by using GPQuest 2.1 to search against a database of 29,188 unique deglycosylated peptide sequences reported in GlycositeAtlas (Sun et al., 2019) and a database containing 252 N-linked glycan compositions. The glycan database was collected from the public database of GlycomeDB (Ranzinger et al., 2011)

Each tandem mass spectrum was first processed in a series of preprocessing procedures, including removing reporter ions, spectrum de-noising, intensity square root transformation (Liu et al., 2007), oxonium ions evaluation, and glycan type prediction (Toghi Eshghi et al., 2016). The top 100 peaks in each preprocessed spectrum were matched to the fragment ion index generated from a peptide sequence database to identify all candidate peptides. All qualified (>6 fragment ions matchings) candidate peptides were compared with the spectrum again to calculate the Morpheus scores (Wenger and Coon, 2013) by considering all the peptide fragments, glycopeptide fragments, and their isotope peaks. The peptide with the highest Morpheus score was then assigned to the spectrum. The mass gap between the assigned peptide and the precursor mass was searched in the glycan database to find the associated glycan. The best hits of all “oxo-spectra” were ranked by the Morpheus score in descending order, in which those with FDR < 1% and covering > 10% total intensity of each tandem spectrum were reserved as qualified identifications. The precursor mass tolerance was set at 10 ppm, and the fragment mass tolerance was 20 ppm.

The quantification of intact glycopeptides was also conducted at the glycoform level. The median log2 ratio value of all the PSMs of an identical intact glycopeptide was used as the relative abundance of the intact glycopeptide. The median relative abundances of intact glycopeptides of samples were normalized to zero. Ratios were then converted to ‘absolute abundance’ scale using estimated abundances in the reference sample.

### Differential expression analysis

In order to distinguish between tumors and non-tumors, we conducted Principal Component Analysis (PCA) on the glycopeptides expressed in all tumor, normal adjacent, and normal ductal samples in the PDAC cohort (Figure 2A). Additionally, we performed Wilcoxon rank sum test analysis on the intact N-linked glycopeptides expressed in tumor and normal adjacent samples from the same cohort. The Benjamini-Hochberg (B-H) method was utilized to correct the p values at 0.01 level of significance to identify significant alternations as potential diagnostic signature glycopeptides (Figure 2B).

To perform gene-annotation enrichment analysis using the KEGG pathway database, GSEApy (Fang, Zhuoqing et al., 2023) was employed on the significantly upregulated and downregulated intact glycopeptides (with FDR < 0.01 and fold change > 2) (Figure 2C). The UniProt knowledge database (UniProt Consortium, 2023) was used to collect cellular location annotations, which were then associated with the gene names of the differentially expressed glycoproteins.

### NMF clustering

We utilized non-negative matrix factorization (NMF)-based clustering to detect the intra-tumor heterogeneity in PDAC. The procedure was conducted similar to our previous work on pancreatic ductal adenocarcinoma (Cao et al., 2021). Briefly, NMF was used to perform unsupervised clustering of tumor samples using the abundances of intact glycopeptides. Only features with CVs in >25% quantile were used for subsequent analysis. The feature matrix was scaled and standardized to z-scores. Since NMF requires a non-negative input matrix, the feature matrix was converted as follows: (1) created one data matrix with all negative numbers zeroed, (2) created another data matrix with all positive numbers zeroed and removed the signs of negative numbers, and (3) concatenated both matrices resulting in a data matrix with positive values and zeros only. The resulting matrix was then subjected to NMF analysis leveraging the NMF R-package (Gaujoux and Seoighe, 2010).

To determine the optimal factorization rank k (i.e., number of clusters), a range of k of 2 to 5 was tested using default settings with 50 iterations. The optimal factorization rank k=3 was selected since the product of cophenetic correlation coefficient and dispersion coefficient of the consensus matrix was high compared to other tested k values. The NMF analysis was repeated using 500 iterations for the optimal factorization rank k. A list of representative features for each cluster was extracted based on the relative basis contribution towards the cluster (threshold was set at 0.8).

### Survival analysis

The survival analysis pipeline in GPnotebook integrated python packages of lifelines (Davidson-Pilon, 2019) and scikit-survival (S. Pölsterl, 2020) to investigate the associations between overall patient survival and intact glycopeptides expression. The Kaplan-Meier curve of overall survival time (Figure 4B) and univariate Cox regression analysis (Figure 4C) were conducted in lifelines to identify the IGPs that were significantly associated with the overall survival time of PDAC cancer patients (adjusted p-value < 0.05). For the three intact glycopeptides with high hazard ratio values, the time-dependent risk analysis was conducted in scikit-survival to estimate the individual trend curves of risk during the time range of tumor development (Figure 4D). To construct a multivariate COX regression model for prognosis, we separated the sample to training set (80% samples, N=68) and testing set (20% samples, N=17). Considering the high dimensional feature space (IGPs size >> sample size) and potential high correlation among certain features, we employed a penalized COX model with elastic net penalty score for feature selection and constructed the COX regression model using 29 independent prognostic IGPs obtained from the training data set. Lastly, we performed the multivariate COX regression model on the testing data and estimate that the model performance using AUC (Figure 4F).

### Glycosylation-phosphorylation cross talk analysis

We investigated the relationship between protein glycosylation and phosphorylation in two conditions: glyco-phospho co-occurrence and correlations. The phosphorylation datasets were collected from cptac package (Lindgren et al., 2021).

In the investigation of glyco-phospho co-occurrence, the overrepresentation analysis was conducted using WebGestalt (Liao et al., 2019) to reveal the KEGG pathways significantly enriched (FDR < 0.05) in the glyco-phospho co-existing proteins (Figure 5B). We applied the linear regression model to demonstrate that the expression values of glycopeptides and phosphopeptides of the identical proteins were positively proportional (p value < 0.01) (Figure 5C).

In order to further investigate the relationship between protein glycosylation and phosphorylation across different proteins, the correlations between protein glycosylation and protein phosphorylation in the CPTAC PDAC tumors (n=105) were studied. The threshold of spearman correlation value > 0.6 was set to filter out the significant correlations between glycopeptides and phosphopeptides. The hub genes (associated with most significant correlations) were highlighted in Figure 5D.

The over-representation analysis was conducted using GSEApy (Fang, Zhuoqing et al., 2023) in the gene names of all phosphopeptides significantly correlated with glycopeptides of BGN at FDR < 0.05 to illustrate the potential phosphorylation regulated signaling pathways impacted by glycosylation alterations.

The protein-protein interaction analysis of the gene names of strongly correlated glycopeptides and phosphopeptides (> 10 positive correlations, spearman corr > 0.6) was performed using API function of STRINGdb (Szklarczyk et al., 2021). GPnotebook integrated the enrichment analysis in the network construction to highlight the pathways involved in the glycoprotein-phosphoprotein cross talk network.

### Biosynthetic glycosylation pathway analysis

We collected 111 glycosylation enzymes involved in the biosynthetic glycosylation pathways. According to the glycosylation steps, the 111 enzymes were classified to three major pathways: precursor, trimming, and capping (Supplementary Table). The spearman correlation between the transcriptomic data and glycoproteomic data were calculated in OmicsOne (Zhang et al., 2021). The differential expression analysis of the global proteomic data was applied on all the cancer cohorts using Wilcoxon rank sum test and set threshold of FDR < 0.01. The spearman correlation values between the enzymes and different glycosylation types were compared using T-test and threshold of p-value < 0.01.

### Software Availability

GPnotebook is readily accessible via https://www.biomarkercenter.org/gpnotebook. Users can effortlessly install the package by executing the command pip install in their preferred shell. Following installation, all notebooks can be executed locally, offering users the flexibility to explore GPnotebook’s functionalities on their own systems.

